# Stochastic Mechanism of Dominant Follicle Selection: Selection of One Suppresses Selection of Others

**DOI:** 10.64898/2026.02.23.707552

**Authors:** Zhuoyan Lyu, Anatoly B. Kolomeisky

**Affiliations:** Department of Chemistry, Rice University, Houston, Texas 77005, USA; Department of Statistics, Rice University, Houston, Texas 77005, USA; Center for Theoretical Biological Physics, Rice University, Houston, Texas 77005, USA; Department of Chemical and Biomolecular Engineering, Rice University, Houston, Texas 77005, USA; Department of Physics and Astronomy, Rice University, Houston, Texas 77005, USA

**Keywords:** ovary, follicle selection, FSH dynamics, stochastic modeling

## Abstract

One of the most critical steps in human reproduction is the selection of the dominant follicle when a single follicle is chosen from a large group of follicles to ovulate. Although this process involves complex hormonal regulation, the complete microscopic picture of unique selectivity remains unclear. We propose a novel stochastic mechanism for dominant follicle selection that incorporates the actions of the most relevant hormones, follicle-stimulating hormone (FSH) and estradiol. Our theoretical picture suggests the following sequence of events. As soon as the FSH concentration reaches the critical threshold, one of the available follicles is randomly selected, which immediately stimulates the production of estradiol, which, via a negative feedback mechanism, suppresses further FSH production, lowering its concentration below the critical threshold. This suppression limits the time window for the possible second follicle selection event, allowing only a single follicle to be selected. Based on this picture, a minimal quantitative theoretical model of dominant follicle selection is developed and analyzed using analytical calculations and computer simulations. Theoretical analysis shows how the interplay between different parameters that govern follicle selection leads to high selectivity. Our theoretical approach can explain some key known observations, providing a quantitative tool for analyzing biological reproduction phenomena.

## 1. Introduction

In sexually mature females, each menstrual cycle typically involves the selection and development of a single follicle that ultimately leads to ovulation [1–5]. This process is known as dominant follicle selection and it is striking and biologically important: it ensures the release of only one egg per cycle, reducing the possibility of multiple pregnancies while optimizing reproductive efficiency by concentrating all resources on the ovulation of the most developed follicle [2, 5–8]. However, the ovary contains many antral (preselected) follicles during each cycle that are, in principle, all capable of responding to hormonal signals and undergoing maturation. Understanding how only one of these follicles is selected, while others are discarded under normal physiological conditions, is a fundamental question in reproductive biology [7, 9–12].

The selection of a dominant follicle is a complex physiological phenomenon that involves multiple biochemical and biophysical processes [13–15]. A central role in these processes is played by follicle-stimulating hormones (FSH) [16, 17], which are produced by the pituitary gland, a small endocrine gland located in the brain. In the early stages of the menstrual cycle, FSH levels begin to increase, promoting the growth and development of the preselected antral follicles, which are capable of maturing and ovulating. The number of preselected follicles generally ranges between 10 and 20 [18, 19]. However, in general, only one follicle, termed the dominant follicle, escapes atresia (follicle death) and continues to grow, while the rest stop developing and die off. A key regulatory component of this process is the negative feedback circuit caused by estrogen hormones, primarily estradiol, which are mostly secreted by the growing selected follicles themselves [20]. As follicles develop under FSH stimulation, they begin to produce estrogen, which in turn suppresses further FSH secretion. This feedback loop, as well as several other hormonal circuits in the ovary [12], is believed to create a self-limiting environment that leads to the selection of only a single dominant follicle [10, 18, 19, 21, 22]. It is important to note here that although choosing a single follicle is the most probable event in this process, sometimes more than one follicle can be selected (although with relatively low probability), which could lead to fraternal twins [1]. It is also possible to have no ovulation events at all, although this is usually a sign of some health problems [23–26]. However, while this qualitative picture of events during follicle selection is widely accepted, the precise microscopic mechanisms of this process remain poorly quantified [12, 21].

The selection of the dominant follicle is also a striking example of unexpectedly high precision in the biological system that being regulated by a large number of random, noisy biochemical processes. Several ideas to explain this phenomenon have been proposed [12, 27–32]. The first approach suggests that preselected follicles at the beginning of the menstrual cycle, are not homogeneous and that they have different sizes and different ranges of various properties, such as the number of FSH receptors [33]. Then, it is argued that the most fitted follicle, probably the one with the largest size or with the largest number of FSH receptors, will become the dominant one. Although this scenario is plausible, it has several problems. Antral follicles (preselected) have a huge number of FSH receptors, and a slight variation in their number should not lead to such dramatic differences. In addition, it is still unclear if the largest follicle by size is chosen for maturation and ovulation [12]. Furthermore, this approach is mostly qualitative. Although there are mathematical models that support the idea that follicles that are more sensitive to FSH are more likely to become dominant, it is not always the largest follicle or the one that is most sensitive to FSH that becomes dominant. Additionally, some of these mathematical approaches are very complicated and involve too many parameters [34, 35].

A different, mathematically elegant approach has also been proposed to explicitly model the effect of various hormones in the dominant follicle selection [12, 30–32, 36]. It argues that the dominant follicle is neither too small and not too large. It was suggested that this might be the result of the biphasic effect of some hormones, namely, that these hormones are active only for intermediate sizes of follicles. However, the predictions of this approach, although physiologically reasonable [12], have also not been supported by data so far. In addition, the biphasic effect has not yet been confirmed. Thus, although the majority of studies indicate that typically the largest follicle becomes dominant, the underlying mechanism remains incompletely understood.

In this study, an alternative theoretical picture of how the selection of the dominant follicle might occur is presented. We hypothesize that there is a stochastic process that explores the effective interactions between FSH and estradiol hormones, creating a narrow temporal “selection window” during which follicles can respond to FSH levels exceeding a critical threshold. This time interval, as we propose, is short enough to allow, on average, only a single follicle to be selected before the hormonal feedback mechanism diminishes the FSH concentration below the selection threshold, effectively closing the gate for other follicles. Our goal here is to present a theoretical method that accounts for the most relevant processes in this system. It allows us to develop a minimal quantitative model of the dominant follicle selection that we are able to analyze analytically and via computer simulations. Our results suggest that the hormonal architecture of the menstrual cycle naturally creates a constraint on the timing and the probability of follicle selection, ensuring that preferentially only one follicle is selected for maturation under typical conditions. Additionally, the model offers insights into scenarios where this process may break down, such as the possibility of more than one follicle being selected under normal conditions, and in polycystic ovary syndrome (PCOS) [25, 26, 37], where either no follicle is selected or multiple follicles compete without a clear dominant winner. Our theoretical method might also be relevant in assisted reproduction, where exogenous hormones may widen the selection window and increase the number of selected follicles.

## 2. Methods

### Theoretical model

Consider a simple theoretical model for the selection of dominant follicles as shown in Fig. 1. We start at the beginning of the menstrual cycle and follow the dynamics of different processes in the system. It will be assumed also that there are *n* (*n* ≃ 10 − 20) antral follicles that can be selected with equal probability for the following maturation and ovulation. The stage before selection is labeled as recruitment in Fig. 1.

**Figure 1.**
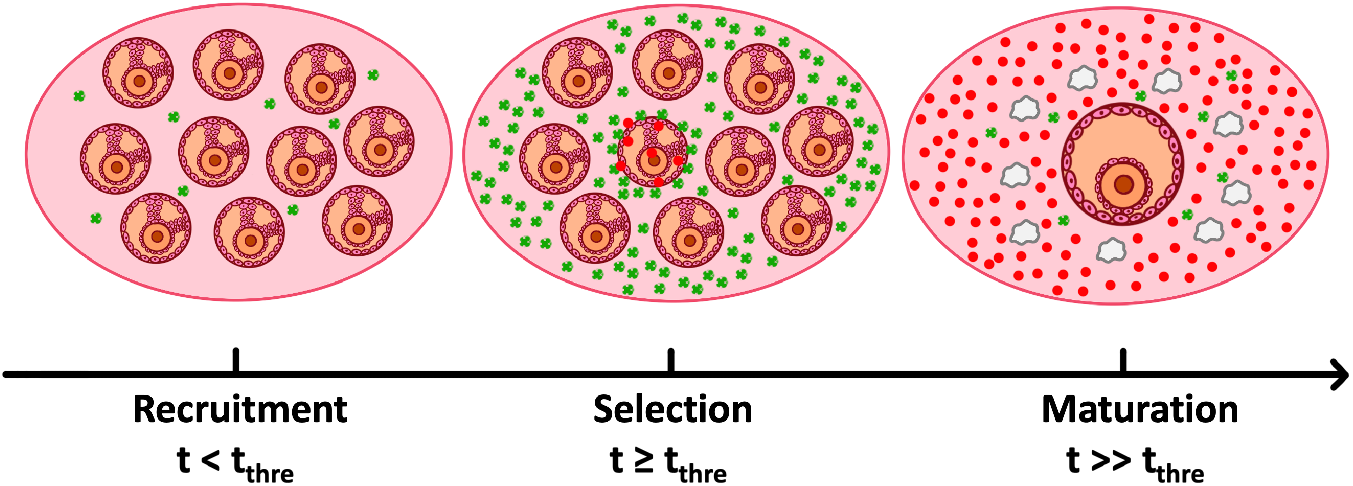
Qualitative overview of the follicle selection process. The green dots represent FSH molecules, and the red dots represent estradiol molecules. During the early period of the menstrual cycle (left panel), the amount of FSH is increasing until the threshold concentration is reached. During the mid-follicular phase, when FSH concentration is higher than the threshold, one follicle is randomly selected among the available follicles. After the selection, it starts to produce estradiol that suppresses the FSH production. During the late follicular phase, FSH concentration drops, the dominant follicle grows into a preovulatory follicle, and the rest of them undergo atresia.

The central role in the selection is played by changes in FSH concentrations defined as *C*(*t*). At *t* = 0, it starts with *C*(0) = *C*_0_. FSH is produced in the pituitary gland in the brain, and it will be assumed that the production rate is *q* while the degradation rate constant for these protein molecules is given by *k*. Then, the dynamics of FSH in the recruitment stage of the selection process can be described as

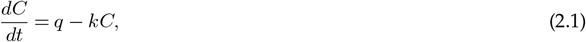

with initial condition *C*(0) = *C*_0_. If the selection process does not occur, the concentration of FSH would follow the dynamics presented by the solution of Eq. (2.1),

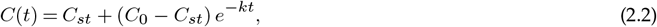

where *C*_*st*_ = *q/k* is the stationary concentration of FSH in the case of no selection of the follicles.

It is known that to select the follicle, the concentration of FSH must rise above some threshold that we define as *C*^∗^ [22]. It will be assumed that, under normal conditions, *C*^∗^ *< C*_*st*_. The FSH concentration reaches the threshold at time *t*^∗^ that can be explicitly estimated from Eq. (2.2),

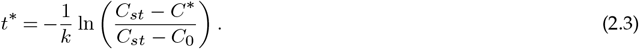

After that time, the follicles can be selected. The main idea of our approach is that this is a stochastic process and that all follicles have the same chance to be selected. After *C*(*t*) reaches *C*^∗^, the selection rate of the first follicle will be assumed to be equal to

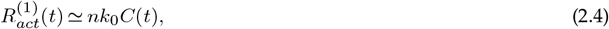

where *k*_0_ is the effective rate constant for selection. This expression can be understood using the following arguments. To select the follicles, the FSH molecules must collide and associate with the follicles. Since there are *n* preselected follicles, the selection rate must be proportional to *n*. In addition, the rate must be a function of the concentration of FSH since the more FSH in the ovary, the more likely for selection to happen. As the simplest approximation, we will assume that 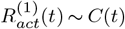 although more complex scenarios can be accounted by extending this approach.

Once a specific follicle is selected at time *t*_1_ ≥ *t*^∗^, it immediately starts to produce a large amount of estrogens that, using the negative feedback mechanism, suppress FSH production. We assume here that estrogen could have been produced even before the selection, but its concentration before the selection is probably too low and can be neglected in our analysis. This leads to the modification of the temporal evolution of FSH concentration after the selection, which can be described as

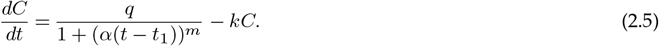

In this expression, *α* is the estrogen feedback strength parameter, which can also be viewed as the effective rate of estrogen production, and *m* is the Hill coefficient that controls the steepness of the feedback response. The Hill coefficient is a phenomenological parameter (*m* ≃ 2 − 4), and it reflects the cooperativity of the feedback regulation events [15, 28, 38, 39]. Eventually, the FSH concentration starts to decrease, and it goes below the threshold concentration at some later time.

The schematic picture of events during the selection of the dominant follicle is presented in Fig. 2. The FSH concentration initially increases (blue curve in Fig. 2) until the threshold concentration *C*^∗^ is reached at time *t*^∗^. Then, at time *t*_1_, one of the *n* preselected follicles is selected. This leads to the production of estrogens that suppress the secretion of FSH, and the concentration profile follows the violet curve (see Fig. 2). At time *t*_*max*_, the FSH concentration reaches its maximal value, and it starts to decrease. After the time interval *Δt* (counting from the moment of first follicle selection), as indicated in Fig. 2, it goes again below the threshold. This time window *Δt* provides the opportunity for other follicles to be selected. The main assumption of our theoretical method is that only a single follicle is selected if the time interval *Δt* is smaller than the average time before the second selection event might occur. The selection rate for the second follicle is given by

**Figure 2.**
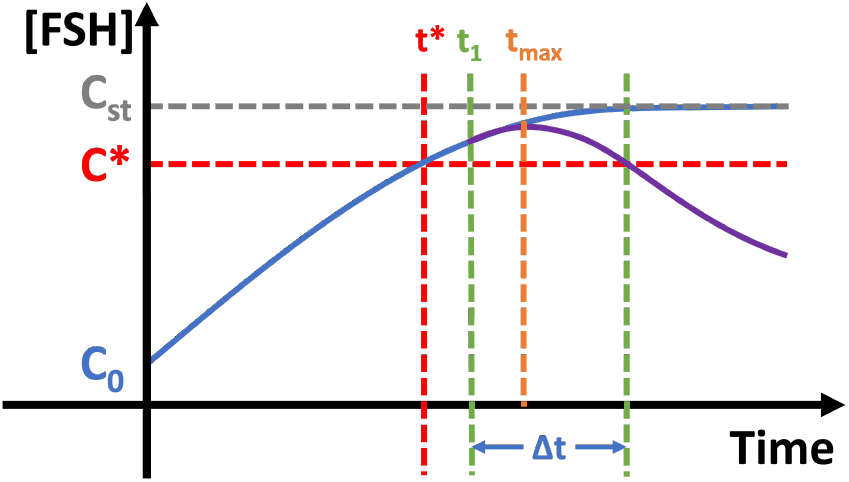
Schematic representation of the FSH concentration evolution and the appearance of a window for follicles selection. The FSH concentration first achieved the threshold *C*^∗^ at time *t*^∗^. The first follicle is selected at time *t*_1_. The concentration profile reaches its maximum at *t*_*max*_. The follicles’ selection might occur only in the time interval *Δt*.

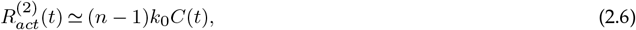

since there is one fewer available antral follicles in the system after activating the first one. Note that the average time mentioned above reflects the stochasticity of the system since this time is averaged over multiple random realizations of this process.

To analyze the conditions that lead to the single dominant follicle, one can use the following approximate analytical arguments. For simplicity, we assume that the selection occurs immediately after passing the threshold, i.e., *t*_1_ ≈ *t*^∗^. Then, FSH is suppressed by estrogens immediately after crossing the threshold, meaning that the FSH concentration does not change much from *C*^∗^. Now we can assume for the maximal FSH concentration, *C*(*t*_*max*_) ≈ *C*^∗^, allowing us to approximate the time at which this occurs (*t*_*max*_ in Fig. 2) from Eq. (2.5),

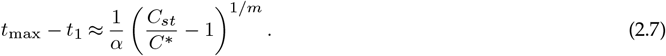

In the next step, we can argue that since the FSH concentration profile does not increase too much in comparison with the threshold *C*^∗^, one can estimate the time interval for the selection window as *Δt* ≈ 2(*t*_*max*_ − *t*_1_), leading to

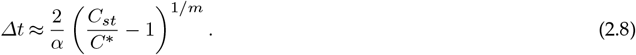

This time interval should be compared with the average time before the second follicle selection might happen, *Δt*_*act*_, which gives

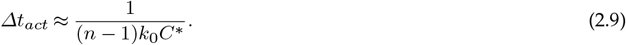

In this expression, we assumed that this happens when *C*(*t*) is not too different from the threshold concentration *C*^∗^, and that there are (*n* − 1) available follicles for the selection. The condition to have only a single dominant follicle is then read as *Δt* ≤ *Δt*_*act*_, which leads to the following relationship,

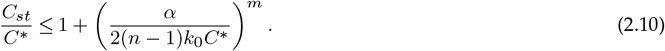

This inequality describes the range of parameters at which the ovary will preferentially have a single dominant follicle for maturation and ovulation. More than one follicle can be selected when this inequality is broken.

Since the analytical analysis presented above involves several strong approximations, we will also employ Monte Carlo computer simulations to test the stochastic model of dominant follicle selection, as detailed in the Supporting Information. More specifically, C++ was used for computer simulations, and python was used for plotting the results.

## 3. Results

### (a) Analysis of Follicles Selection Processes Using Approximate Analytical Approach

Our theoretical method allows us to fully evaluate the temporal evolution of the FSH concentration using analytical solutions from Eq. (2.2) and numerical solutions of Eq. (2.5). Assuming that the follicle selection occurs immediately after the FSH concentration reaches the threshold, the dynamics of the FSH concentration changes with time is presented in Fig. 3. The chosen parameters ensure that the obtained time scales are consistent with known information that the dominant follicle selection occurs around days 5-7 after the beginning of the menstrual cycle [1]. As one can see, Fig. 3 reproduces well the behavior qualitatively predicted above (compare with Fig. 2), and this should allow us to explicitly evaluate the time interval during which the follicle selection might occur. Using the parameters utilized in Fig. 3, we estimate that the time window is around *Δt* ≈ 0.5 day. This relatively small time interval should not generally allow another selection of the follicle, supporting the high selectivity of the process. Our theoretical results are also consistent with experimental observations [40], showing that FSH peak concentration (around day 5) aligns with the time when one follicle becomes dominant.

**Figure 3.**
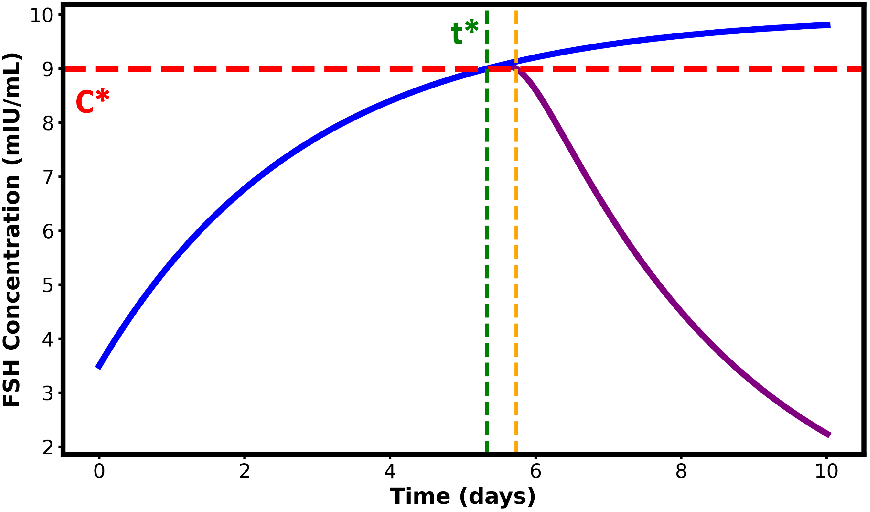
Temporal evolution of FSH concentration assuming that the first follicle selection occurs immediately after the threshold concentration is achieved. The blue line is the dynamics of FSH when selection process doesn’t occur; the purple line is the dynamics of FSH when selection process occurs at *C*^∗^; the red dashed line is the FSH concentration threshold *C*^∗^; the green dashed line is the time of selection (*t*_1_ = *t*^∗^); the orange dashed line is the time when FSH concentration drops below the threshold *C*^∗^. Parameters used for calculations are: the FSH stationary concentration if no selection occurs is *C*_*st*_ = 10 mIU/ml; the threshold concentration is *C*^∗^ = 9 mIU/ml; the estrogen production rate after the follicle selection is *α* = 2 day^−1^, and the Hill coefficient is *m* = 3.

The results in Fig. 3 also indicate that during the time window of follicle selection the FSH concentration does not deviate much from the threshold *C*^∗^, allowing us to apply the analytical framework presented above. More specifically, Eq. (2.10) can be used to investigate the ranges of parameters at which the selection of only a single follicle might occur. The results of our analysis are presented in Fig. 4.

**Figure 4.**
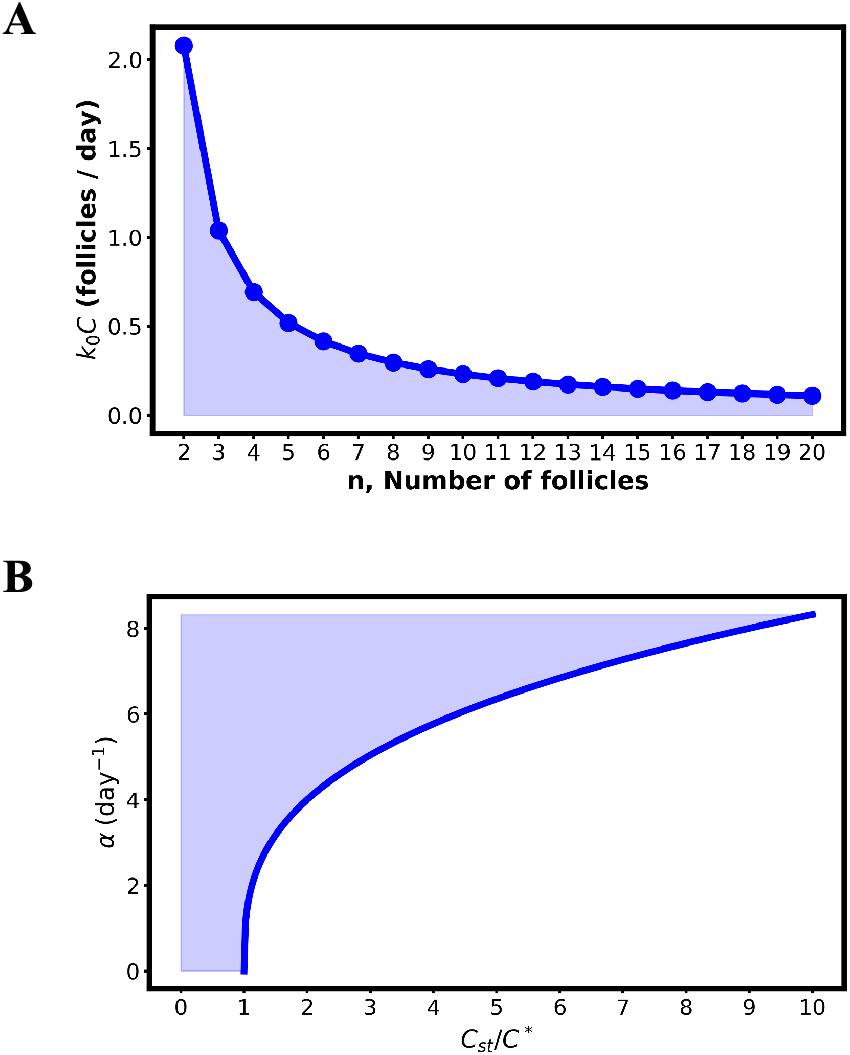
Boundaries for separating the dynamic regimes of selection of single follicle versus the selection of more than one follicle: (A) by varying the number of preselected follicles *n* and the effective rate of selection per one follicle, *k*_0_*C*^∗^; (B) by varying the estrogen production rate *α* and the ratio of stationary and threshold concentrations, *C*_*st*_/*C*^∗^. Blue shaded areas correspond to the regions where only a single follicle is selected. The following parameters were used in calculations: (A) 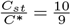, *α* = 2, *m* = 3, (B) *m* = 3, *n* = 10, 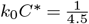.

Fig. 4A shows the boundary for the single follicle selection when the number of preselected follicles *n* and the effective rate of selection per one follicle, *k*_0_*C*^∗^, are varied while the other parameters of the system are kept fixed. One can see that if the number of preselected follicles is relatively small, even for large selection rates, only a single follicle will typically be selected. However, increasing the number of follicles will require much slower selection rates to keep the high selectivity of the dominant follicle selection. This observation might explain why the number of antral follicles is not too large at the beginning of the menstrual cycle, despite the ovary having a very large number of follicles. Our theoretical method suggests that for large *n* it will be difficult to support the selection of only one follicle since a very limited range of selection rates can support high selectivity.

Fig. 4B shows the boundary for the single versus multiple follicles selection when the estrogen production rate *α* and the ratio of stationary and threshold concentrations, *C*_*st*_/*C*^∗^, are varied while other parameters are fixed. In this case, when the threshold concentration is close to the stationary concentration (*C*_*st*_/*C*^∗^ ∼ 1), to select only a single follicle would be possible for a large range of estrogen production rates, including relatively weak rates. However, increasing this ratio (*C*_*st*_/*C*^∗^ ≫ 1) would still produce the high selectivity for a smaller range of faster estrogen production rates. This can be easily understood since for larger *C*_*st*_/*C*^∗^ the time interval of selection would increase unless more estrogen is produced to suppress the FSH concentration sooner.

### (b) Stochastic Simulations of Follicle Selection Processes

While our theoretical approach provides an analytical description of processes taking place in the ovary during the follicle selection, it relies on several strong approximations that might not always work. They include the assumption of immediate follicle selection after the FSH concentration reaches the threshold and the assumption that the concentrations of FSH during the time window for selection are not too different from the threshold *C*^∗^. To test the approximate analytical framework and to get a better understanding of underlying processes, we performed stochastic Monte Carlo computer simulations. The details of the simulations as well as the parameters used in our calculations are explained in the Supporting Information.

The results of our computer simulations, using realistic parameter sets, are presented in Figs. 5 and 6. First, we analyzed the probability of having different numbers of follicle selections during the selection time window (see Fig. 5A). It is shown that under normal conditions, more than 90% of events exhibit a single follicle selection, while in less than 10% of cases, two follicles are selected. We have not observed the selection of more than two follicles in our simulations. Thus, our theoretical method suggests that selecting a single dominant follicle is a preferred dynamic behavior, but due to stochasticity, the chance of activating more than one follicle is small but not zero. This naturally explains the observations in healthy normal females that typically exhibit maturation and ovulation of only a single follicle per menstrual cycle [1].

**Figure 5.**
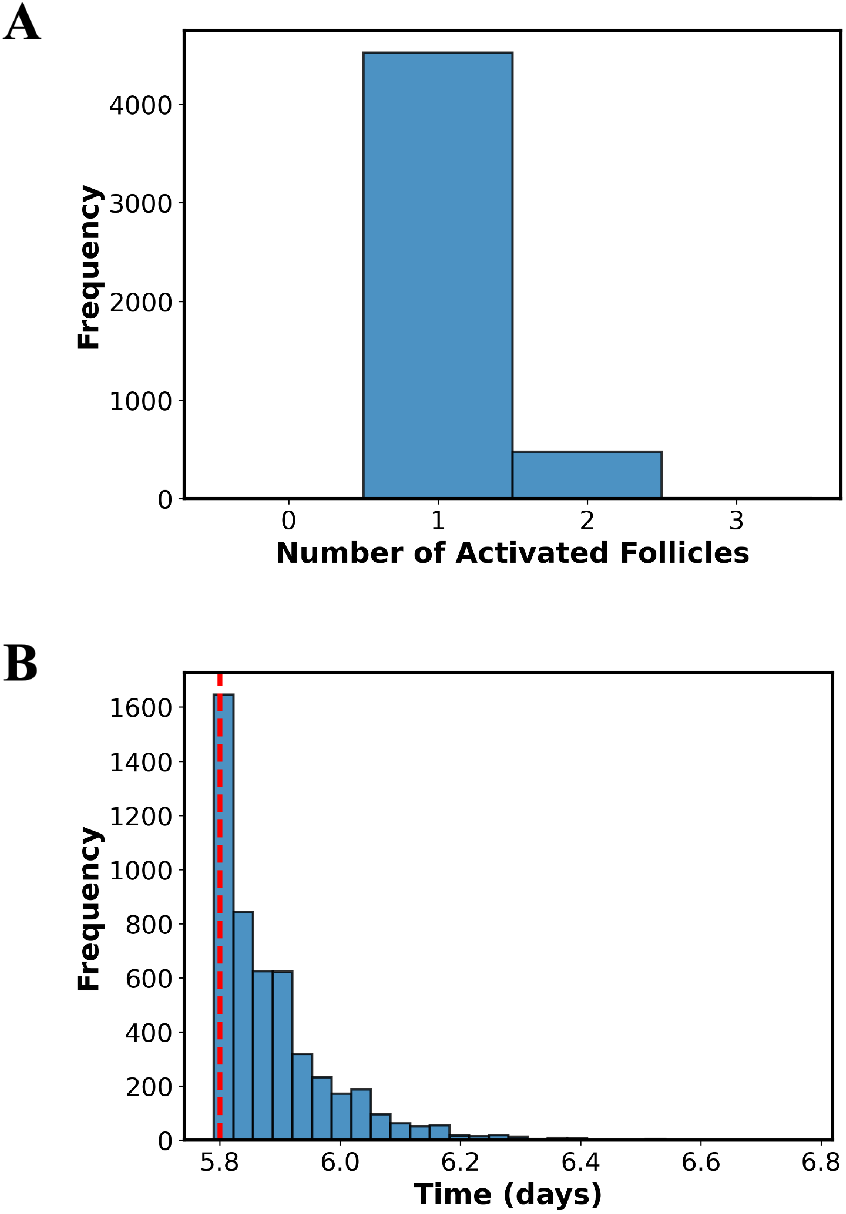
Simulation results for the follicle selection processes. (A) The frequency of activating different numbers of follicles (after 5000 simulations). (B) The distribution of times when the first selection occurred. The red dashed line is the time when FSH concentration reaches the threshold. The following parameters were used in simulations: the initial FSH concentration *C*_0_ = 3.5 mIU/ml, the FSH concentration threshold *C*^∗^ = 9.8 mIU/ml, the FSH production rate *q* = 6.0 mIU/ml × day^−1^, the degradation rate *k* = 0.6 day^−1^, the estrogen production rate *α* = 100 day^−1^, the Hill coefficient *m* = 3, and the number of preselected follicles *n* = 10.

**Figure 6.**
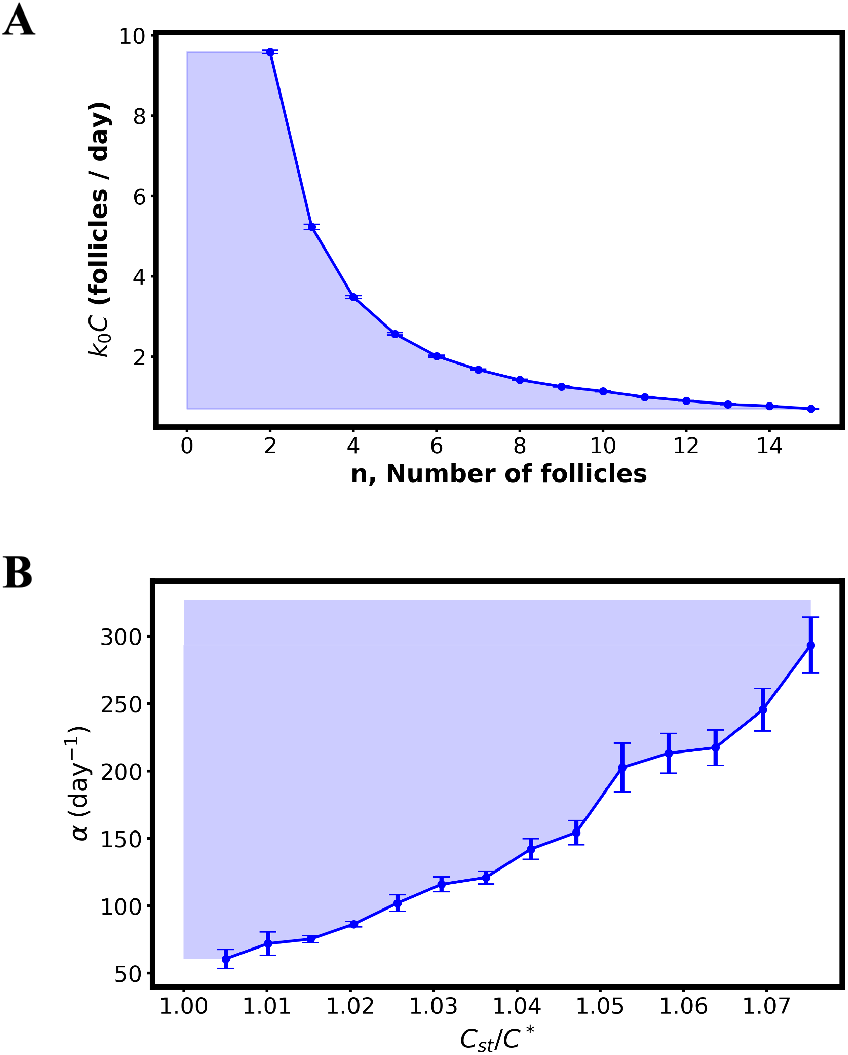
(A) Simulation result for the maximum allowable value of *k*_0_ × *C* for each number of follicles *n* when the probability of having more than 1 follicle being selected is less than 10%. Parameters used in the simulation are: *C*^∗^ = 9.8 mIU/ml, *q* = 6.0 mIU/ml × day^−1^, *k* = 0.6 day^−1^, *α* = 100, *m* = 3, and *C*_0_ = 3.5 mIU/ml. (B) Simulation result for the minimum allowable value of estrogen feedback rate *α* and *C*_*st*_/*C*^∗^ when the probability of having more than 1 follicle being selected is less than 10%. Parameters used in the simulation are: *C*_*st*_ = 10 mIU/ml, *q* = 6.0 mIU/ml × day^−1^, *k* = 0.6 day^−1^, *m* = 3, *k*_0_ = 0.1 day^−1^, and *C*_0_ = 3.5 mIU/ml, with *n* = 10 total follicles. Error bars are SEM (standard error of the mean).

Fig. 5B presents the computer simulations for the distribution of times before the selection of the first follicle occurs. As one can see, it does not happen immediately after the FSH concentration reaches the threshold *C*^∗^ (red vertical dashed line in Fig. 5B). At the same time, the distribution is relatively narrow (less than 0.5 of a day for almost all events), and the significant fraction of selection events do happen quickly after passing the threshold. These results qualitatively support the approximations that we utilized in our analytical calculations.

Using computer simulations, we also investigated the range of parameters that support the selection of only one follicle. Our computations suggest that the probability of activating two follicles is less than 10% for typical conditions in healthy females (see Fig. 5A). This is a realistic estimate, and it can be explained using the following arguments. The production of two follicles for the ovulation can lead to the so-called fraternal twins [1, 12]. The probability of such events is known to be ∼ 1 − 3% [41]. However, only 10-20% of twin eggs obtained in such events survive [42], and this means that the probability of activating more than one follicle should be less than 10%, in agreement with our estimate. This is the value that we use in our simulations (see Fig. 6) as the borderline to identify the ranges of parameters that lead to single dominant follicle selection versus multiple follicle selection.

The results presented in Fig. 6 qualitatively agree with the predictions from the approximate analytical theory (compare with Fig. 4). If the number of preselected follicles is relatively small, then the single follicle can be chosen for the maturation for large range of selection rates, even if they are relatively slow (Fig. 6A). However, increasing the number of antral follicles (*n*) lowers the range of selection rates that can lead to the highest selectivity: see Fig. 6A. In this case, only fast production of the estrogen can lead to the single follicle selection. Note also that the sensitivity of choosing the proper selection rate is relatively small for *n* ≃ 10 − 20. This means that slight variations in the number of preselected follicles, which are realistic, will not affect the high selectivity. In addition, as shown in Fig. 6B, if the ratio of the FSH stationary and threshold concentrations is small and close to unity, a large range of estrogen production rates can achieve the selection of only one follicle, including the slow rates. But increasing the ratio forces the system to have a shorter range of generally faster estrogen production rates that can sustain the single follicle selection.

## 4. Summary and Conclusions

In this paper, we presented a novel theoretical approach to explain the dominant follicle selection processes in females’ ovaries during each menstrual cycle. The main idea of our method is that several stochastic processes can lead to unusually high selectivity of follicle selection. According to our approach, the following microscopic events are taking place in the system. Starting from the beginning of the menstrual cycle, the FSH concentration starts to increase. Around days 5-7, it reaches the critical threshold after which the selection becomes possible. We view the follicle selection as a random stochastic process that depends on the FSH concentration and on the number of preselected follicles. As soon as the first follicle is selected, it begins to produce estrogens that suppress the production of FSH molecules. This leads to a decrease in the FSH concentration that soon passes the threshold level again. The time interval when the FSH concentration is above the threshold and the first follicle is selected determines the window of opportunities for the second follicle selection, which is also a stochastic event that depends on the number of available follicles and current FSH concentration. Since such selections are random, if this time interval is smaller than the average time before the second follicle can be selected, this leads to choosing only a single dominant follicle. However, because of the stochastic nature of these processes, sometimes more than one follicle can be selected in this process. It is important to note here that the existence of a selection window has been widely accepted. However, the novelty of our approach is that we view all the preselected follicles the same and time interval is the only stochastic quantity that can be tuned by hormonal circuits to make sure that after the first selection, which also occurs at a random time, the next stochastic selection event, on average, does not occur.

Based on these arguments, we developed a minimal quantitative theoretical model that took into account the activities of the two probably most relevant players in these processes, FSH and estrogen hormones [1–4]. The model was solved analytically under some approximations. It was also investigated using Monte Carlo computer simulations. Our theoretical calculations identified the range of parameters at which the single follicle is preferably selected for maturation and ovulation. It is also shown that, under normal conditions, more than 90% of the events should lead to the selection of a single follicle, while less than 10% might produce more than one dominant follicle. Our theoretical framework can also explain the relatively small number of preselected follicles. It is argued that having too many antral follicles would not make the system robust since it will not easily reach the high selectivity, which is a preferred mode of action in ovulation. In addition, it is suggested that the single follicle selectivity requires the threshold concentration to be close enough to the stationary concentration, and this is also related to the robustness of the system. Some aspects of our theoretical approach can be tested in experiments since it provides quantitative predictions of the probability of single follicle selection on the number of preselected follicles, on the ratio of critical concentration to the stationary FSH concentration, and on the rate of estrogen production.

The proposed theoretical framework is consistent with various observed patterns of follicle selection in natural menstrual cycles. It correctly predicts that the great majority of such events in healthy females lead to a single dominant follicle; however, there is a small chance of having more than one dominant follicle, which is a result of the stochasticity of underlying microscopic processes. We can also speculate about the observations of increased probability of multiple selected follicles in older females [43] by suggesting that this could be due to an increase in FSH concentration which triggers a lowering of the estrogen production rate and/or due to a change in the stationary and threshold concentrations of FSH (see Figs. 4B and 6B). Importantly, our theoretical approach can also explain other outcomes in the selection process. For example, conditions such as polycystic ovary syndrome (PCOS) that could lead to zero selection of follicles [25, 26]. In our theoretical method, this might happen if the FSH threshold concentration is larger than the stationary concentration (*C*_*st*_/*C*^∗^ *<* 1), yielding no conditions under which the selection of any follicle can happen. Similarly, in assisted reproductive treatments, administering exogenous FSH may raise the stationary concentration *C*_*st*_ enough to significantly widen the selection window, intentionally increasing the possibility of multi-follicular recruitment. Our method also suggests other ways to increase the probability of multiple follicle selection by decreasing the estrogen production rate or increasing the ratio of concentrations (also shown explicitly in Fig. S1).

Although the proposed theoretical framework provides physically clear and simple explanations of dominant follicle selection processes, it is rather a very oversimplified approach that neglects many crucial features of these complex phenomena. It is important to discuss the limitations of our theoretical method. Our results have the same order of magnitude of relevant hormones concentrations as experimentally measured data [40]. However, one should notice that the proposed theoretical method mostly focuses on biological trends instead of trying to exactly reproduce the absolute values of hormone concentrations. As hormone level measurements vary across laboratories, the values selected for this study are those that seems to be the most representative, as explained in the Supporting Information. Even though estrogen is already produced by small antral follicles before the dominant follicle appears, those produced amounts are relatively small and can be neglected in the minimal theoretical approach [20]. Additionally, it is possible for dominant follicles to emerge in the early follicular or luteal phase despite lower levels of FSH, but since they typically do not ovulate, they are not considered as “selected follicle” in our theoretical model. Moreover, during the menstrual cycle, many other hormones are typically produced, such as luteinizing hormones, androgens (e.g., testosterone), inhibin hormones, and they also play an important role in follicle selection [6–8, 12]. In addition, our theoretical method neglects the growth of antral follicles that is occurring at the same time [12]. This should definitely influence the various parameters in the model, such as selection and suppression rates. Furthermore, in our analysis, the simplest negative feedback mechanism is assumed, while more complex mechanisms might strongly affect the dynamics of follicle selection. One more important question is, what is the origin of the FSH threshold concentration [44]? However, despite these limitations, the proposed theoretical approach might be utilized as a convenient quantitative tool that can help clarify the microscopic mechanisms of biological reproduction phenomena, which should eventually lead to improvements in females’ health.

## Ethics

This work did not require ethical approval from a human subject or animal welfare committee.

## Data Accessibility

All data and code are available from [45]. The data and code that support the findings of this study have been deposited in the Zenodo repository with the identifier https://doi.org/10.5281/zenodo.17094488. Supplementary material is available online.

## Declaration of AI use

We have not used AI-assisted technologies in creating this article.

## Authors’ Contributions

Z.L. and A.B.K. designed research; Z.L. performed research; Z.L and A.B.K. analyzed data; and Z.L. and A.B.K. wrote the paper.

## Acknowledgements

This work was also performed in part at the Aspen Center for Physics, which is supported by National Science Foundation grant PHY-2210452.

