## Supplemental file for "Stochastic Mechanism of Dominant Follicle Selection: Selection of One Suppresses Selection of Others"

INTERFACE

rsif.royalsocietypublishing.org

Research

Article submitted to journal

**Author for correspondence:**
Corresponding author


### Supporting Information: Stochastic Mechanism of Dominant Follicle Selection: Selection of One Suppresses Selection of Others

Zhuoyan Lyu[1,2], Anatoly B. Kolomeisky[1,3,4,5]

[1]Department of Chemistry, Rice University, Houston, Texas 77005, USA
[2]Department of Statistics, Rice University, Houston, Texas 77005, USA
[3]Center for Theoretical Biological Physics, Rice University, Houston, Texas 77005, USA
[4]Department of Chemical and Biomolecular Engineering, Rice University, Houston, Texas 77005, USA
[5]Department of Physics and Astronomy, Rice University, Houston, Texas 77005, USA

THE ROYAL SOCIETY
PUBLISHING

© The Authors. Published by the Royal Society under the terms of the Creative Commons Attribution License http://creativecommons.org/licenses/by/4.0/, which permits unrestricted use, provided the original author and source are credited.

### 1. Simulation algorithm

We simulated follicular selections using Monte Carlo simulation. The model depends on changing FSH over time and the number of follicles that are still available to be selected. Before any selection event, FSH follows an increasing equation,

$$\frac{dC}{dt} = q - kC.$$

After the first selection, The production of estradiol reduces FSH production according to a Hill function,

$$\frac{dC}{dt} = \frac{q}{1 + [\alpha(t - t_1)]^m} - kC,$$

where $t_1$ is the selection time. Trajectories are integrated by forward Euler with fixed step $\Delta t$ up to a maximum time $t_{\max} = 13$.

Stochastic selection events are generated by inverse-transform sampling for an inhomogeneous Poisson process whose instantaneous rate is

$$R_{act}^{(1)}(t) \simeq nk_0C(t), \tag{1.1}$$

For each possible event, we use $r \sim U(0, 1)$ to set a target cumulative hazard $H^* = -\ln r$. We then increase the time in steps $\Delta t$, accumulating the increment in the discrete $R_{act}^{(1)}(t)\Delta t$ while simultaneously updating $C(t)$ based on the previous dynamics; when the accumulated hazard first exceeds $H^*$, the event time is accepted.

An event is recorded as an selection only if the FSH concentration exceeds a fixed threshold, $C(t) > C^* = 9.8$. After each selection, $n$ decreases by one; if this is the first selection, we enter the post-selection dynamics (setting $t_1 = t$). The loop continues until $t_{\max}$ or $n = 0$.

For fixed follicle count $n$ and a given $k_0$, we perform $n_{\text{sim}} = 1000$ independent realizations and estimate the proportion of simulations with at least two selections. To determine the largest $k_0$ that satisfy a specific condition (here, $\leq 10\%$ with $\geq 2$ selections), we use bisection over a bounded interval $[1e - 4, 1.0]$. If the criterion is already violated at $1e - 4$, no feasible value is reported; otherwise, we iteratively tighten the interval, evaluating the $\geq 2$-selection proportion at the midpoint and moving the lower (upper) bound when the criterion is satisfied (violated). We record 50 maximum $k_0$ values and its achieved proportion.

This procedure is repeated across follicle counts $n = 2, \ldots, 15$, with multiple independent replicates per $n$ obtained by varying the RNG seed. We record 50 maximal feasible $k_0$ values for each $n$.

The same algorithm is applied to the figure for the minimum allowable value of estrogen feedback rate $\alpha$ and $C_{st}/C^*$ when the probability of having more than 1 follicle being selected is less than 10%

### Supplementary Table

| Parameters | Values | Reasons |
|---|---|---|
| $\alpha$ | $100\ \text{day}^{-1}$ | To narrow the selection window. |
| $n$ | 10 | See references [1, 2] |
| $q$ | $6.0\ \text{mIU mL}^{-1}\ \text{day}^{-1}$ | To allow the selection to happen between days 5 and 7: see references [3]. |
| $k$ | $0.6\ \text{day}^{-1}$ | To allow the selection to happen between days 5 and 7: see reference [3] |
| $k_0$ | $0.1\ \text{day}^{-1}$ | To control the number of selected follicles. |
| $m$ | 3 | To narrow the selection window. |
| $C_0$ | $3.5\ \text{mIU mL}^{-1}$ | see reference [4]. |
| $C_{st}\ (= q/k)$ | 10 | See reference [5]. |
| $C^*$ | $9.8\ \text{mIU mL}^{-1}$ | To narrow the selection window. |

**Table 1.** Parameters utilized in calculations and reasons for choosing them.

### Supplementary Figures

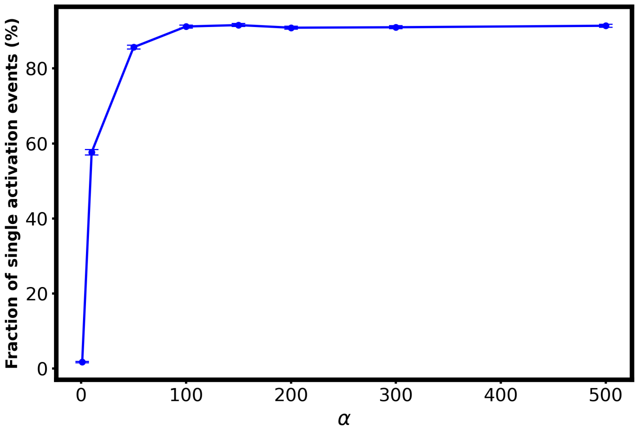


**Figure 1.** Simulation result for the fraction of single selection events for each estrogen production rate $\alpha$. Error bars are SEM (standard error of the mean)

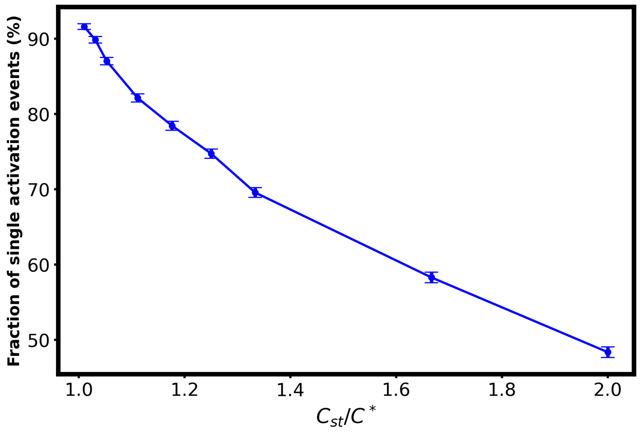


**Figure 2.** Simulation result for the fraction of single selection events for $C_{st}/C^*$. Error bars are SEM (standard error of the mean)

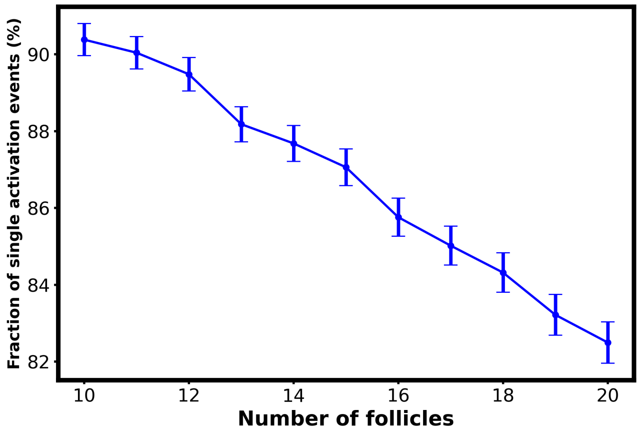


**Figure 3.** Simulation result for the fraction of single selection events for each number of preselected follicles $n$. Error bars are SEM (standard error of the mean)
